# Resolving the data asynchronicity in high-speed atomic force microscopy measurement via the Kalman Smoother

**DOI:** 10.1101/2020.08.10.242719

**Authors:** Shintaroh Kubo, Suguru Kato, Kazuyuki Nakamura, Noriyuki Kodera, Shoji Takada

## Abstract

High-speed atomic force microscopy (HS-AFM) is a scanning probe microscopy that can capture structural dynamics of biomolecules in real time at single molecule level near physiological condition. Albeit much improvement of the instruments, while scanning one frame of HS-AFM movies, biomolecules often change their conformations largely. Thus, the obtained frame images can be hampered by the time-difference, the asynchronicity, in the data acquisition. Here, to resolve this data asynchronicity in the HS-AFM movie, we developed Kalman filter and smoother methods, some of the sequential Bayesian filtering approaches. The Kalman filter/smoother methods use alternative steps of a short time-propagation by a linear dynamical system and a correction by the likelihood of AFM data acquired pixel by pixel. We first tested the method using a toy model of a diffusing cone, showing that the Kalman smoother method outperforms to reproduce the ground-truth movie, compared to that mimics the raw AFM movie, and the Kalman filter result. We then applied the Kalman smoother to a synthetic movie for conformational change dynamics of a motor protein, i.e., dynein, confirming the superiority of the Kalman smoother. Finally, we applied the Kalman smoother to two real HS-AFM movies, FlhAc and centralspindlin, reducing distortion and noise in the AFM movies. The method is general and can be applied to any HS-AFM movies.

## Introduction

Observing biomolecules at work has long been a major challenge in molecular biology, biochemistry, and biophysics. High spatial resolution, i.e., atomic resolution, can be reached by several technologies in structural biology, such as X-ray crystallography, nuclear magnetic resonance, and cryo-electron microscopy (1, 2). Especially, the recent revolution in cryo-electron microscopy technology markedly broadened the range of applicability to obtain atomic resolution information (2). Yet, these methods do not usually observe real-time motions of biomolecules, with some notable exceptions (3, 4). On the other hand, high temporal resolution can be achieved by different types of experiments; for example, laser chemistry offers broad-range of spectroscopy with ultra-high time resolution up to femtoseconds (5, 6). Yet, most these methods do not provide global and structurally resolved information. Thus, reaching high resolution simultaneously in space and time is extremely challenging.

In this respect, the high-speed atomic force microscopy (HS-AFM) has a unique feature of a medium-range spatiotemporal resolution, enabling it to observe single biomolecular structure at work as a video/movie (7–12). The AFM, in general, is a scanning probe microscopy where a needle-like probe attached at the tip of cantilever scans an area of surface (*xy*-plane) (Figure 1A). While scanning, the probe moves up and down (*z*-direction) detecting surface atoms or molecules bound on the surface. Armed by recent technology developments, the HS-AFM enables to scan an area of surface within < 100 ms with a spatial resolution of ~1 nm in *xy*-direction and ~ 0.1 nm in *z*-direction (10). With this spatiotemporal resolution, the HS-AFM successfully observed structural dynamics of many proteins at work, such as the walking motion of myosin V, the rotary motion of F_1_-ATPase, the power-stroke motion of axial dynein, and the target recognition and DNA cleavage of CRISPR-Cas9 (13–16).

**Figure 1.**
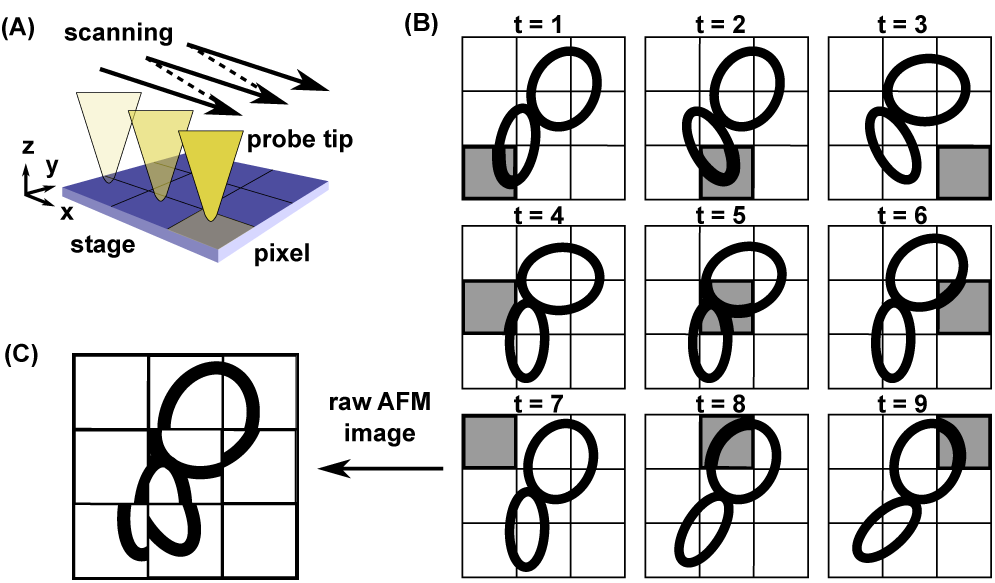
Data asynchronicity in the HS-AFM measurement. (A) In the HS-AFM, the probe tip scans the surface with the bound molecule in the raster mode and sequentially records the molecular height. (B) A “molecule” (cartoon drawn by two ovals) bound on the surface changes its shape along time, t = 1, 2, 3,,,. The HS-AFM observes the molecular height one pixel at a time (grey pixel). (C) A raw AFM image obtained after one complete scan contains data from different times and can be distorted.

While attractive, the HS-AFM has many limitations as well, of which we focus on one in this study. The AFM is the scanning microscopy where the probe tip scans the surface on which target molecules are bound. Typically, it takes ~10 µs to record the data at one pixel. Then, scanning one frame containing, for example, 100 x 100 pixels results in ~100 ms. Therefore, the HS-AFM image obtained after one scan is made of data collected at different time. During this period, target biomolecules can move significantly (Figure 1B). This data asynchronicity can distort the obtained AFM image significantly, depending on the time scale of molecular motions relative to the scanning time (Figure 1C). The apparent image with the data asynchronicity could miss or duplicate a part of molecule, and provide shrunk or stretched shape of molecules(17).

Our purpose in this study is to resolve such data asynchronicity issue in HS-AFM using a sequential Bayesian data assimilation approach (18, 19). The sequential Bayesian data assimilation couples the dynamical system model and the measurement in a sequential manner. The dynamical system model provides an approximate method to propagate the probability distribution of the state of interest for a short time, called “prediction” (Figure 2A. Details are described later). The observation provides information for the state of interest at a time and modifies the probability distribution of the predicted state, termed “filtering”. In the prediction step, we update the probability distribution of the state for the interval between two consecutive observations. In the subsequent filtering process, we modify the probability using the observation data via the Bayesian formula. This is followed by the next round of the prediction and the filtering. Through this iterative process, we can integrate dynamical system information with observation.

**Figure 2.**
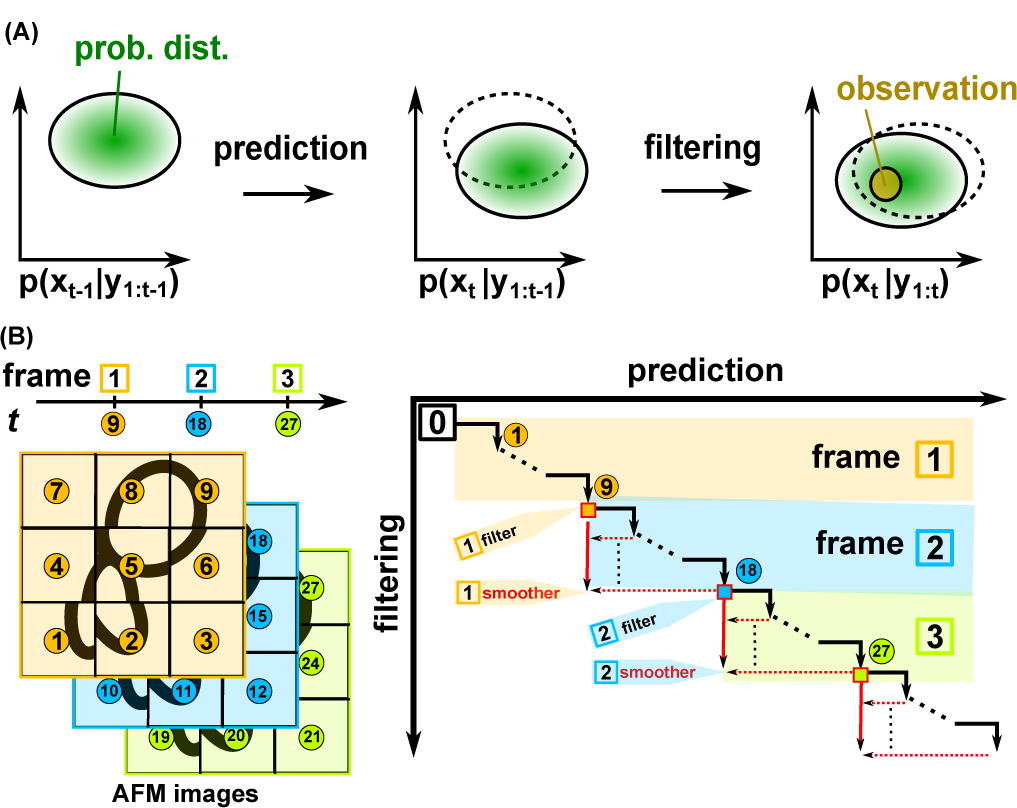
Concept of sequential Bayesian data assimilation and the realization of Kalman filter/smoother in the HS-AFM measurement. (A) Starting with the distribution, *p*(*x*_*t*-1_|*y*_1:*t*-1_), (left), we propagate the state *x* for a unit time via the linear dynamical system model and obtain the distribution, *p*(*x*_t_|*y*_1:*t*-1_), (center) which is called “prediction”. Then, taking into accounts of a new observation *y*_*t*_ at time t, we update the probability distribution, *p*(*x*_t_|*y*_1:*t*_), (right) which is termed “filtering”. The pair of prediction and filtering can be iteratively applied to update the time. (B) Illustration of the method with a 9-pixel toy image. In the Kalman filter approach for HS-AFM data, each measurement corresponds to obtain the molecular height at one pixel. For example, the Kalman filter obtains the first and second frame images using data up to 9 and 18 time points, respectively. The repeat of the prediction and the filtering for 9 times provides the minimal data for one frame (black arrows with circled numbers). In the Kalman smoother for HS-AFM, we also utilize 9 extra-time points in future measurement (red arrows). For example, the Kalman smoother obtains the first and second frame images using data up to t = 18, and 27, respectively.

In the current context of HS-AFM measurement, the state of interest corresponds to an AFM-like image that can be stacked into one-dimensional vector. A single observation corresponds to a data on the molecular height at one pixel. For the sake of generality, we do not assume any specific motions of target molecules. To this end, we employ a widely-used and probably the simplest dynamical system model, the linear dynamical system (LDS). The LDS provides simple and analytical formula for the iterative updates, which is normally called the Kalman filter. The Kalman filter method integrates the observations at the current time as well as the past time points with the LDS, making it applicable as the on-the-fly filter. When the whole time-series observation data are available, i.e., the off-line mode, we can extend the method to utilize the observations in future, as well as the current and past observation data, which is called the Kalman smoother, or the Rauch-Tung-Striebel (RTS) smoother (20).

In this paper, we begin with a brief description of the sequential Bayesian data assimilation approach, the Kalman filter/smoother methods, and their realizations for the HS-AFM data. We then examine them with a twin-experiment for a toy model of a diffusing cone. We show that the Kalman smoother method outperforms the raw AFM movie as well as the Kalman filter method. As a second test, we setup another twin-experiment using a molecular dynamics simulation trajectory of a motor protein, dynein. In the molecular dynamics simulation, dynein exhibits a rapid conformational change, called power-stroke. How this rapid power-stroke can be captured is a question of interest. While the power-stroke was hardly detected by a mimic of raw HS-AFM data, the Kalman smoother could provide, albeit weak, signal of power-stroke. Finally, we applied the Kalman smoother to two real HS-AFM movies, FlhAc and centralspindlin, demonstrating our method to produce molecular envelope shapes. From the Kalman smoother image, we could model possible configurations of the two molecules.

## Methods

### The linear dynamical systems(19–21)

We begin with a stochastic dynamics system described by a time-dependent state-space vector *x*_*t*_ ∈ R^*k*^, where the time *t* is discretized as *t* = 0, 1, 2, and the associated measurement *y*_*t*_ ∈ R^*l*^ that depends on *x*_*t*_. The discrete time corresponds to the time at which new measurement data is acquired. *k* and *l*, respectively, represent the dimension of state-space and the measurement. For the HS-AFM data, *x*_*t*_ corresponds to an AFM-like image of interest, while *y*_*t*_ is the real AFM measurement data at one pixel at a time.

For the state vector, we define the system model of time-propagation,

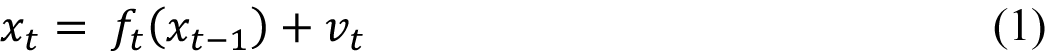

where the function *f*_*t*_ defines the time-propagation and *v*_*t*_ ∈ R^*k*^ represents the system noise. For the Markovian process, this short time propagation can be expressed as *p*(*x*_*t*_|*x*_*t*-1_). We define the measurement system,

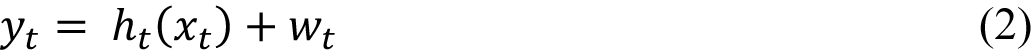

where *h*_*t*_ defines the measurement function, and *w*_*t*_ ∈ R^*l*^ represents the measurement noise.

When the system model is linear and the noise is Gaussian and white, we can write the Eq. (1) as

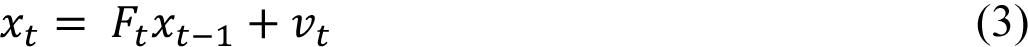

where *F*_*t*_ is a *k* × *k* matrix that define the deterministic part of the system model. The *v*_*t*_ is the Gaussian white noise with its mean vector zero and variance-covariance matrix *Q*_*t*_ (*k* × *k* matrix). We further assume the linear measurement function and the Gaussian white noise in the measurement (2) as

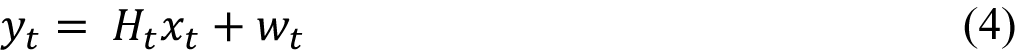

where *H*_*t*_ is a *l* × *k* matrix, the *w*_*t*_ is Gaussian white noise with its mean vector zero and variance-covariance matrix R_*t*_ (*l* × *l* matrix). The equations (3) and (4) constitute the linear dynamical system (LDS).

### The Kalman filter method

In the sequential Bayesian statistical approach, we alternatively take prediction and filtering steps(22). The prediction step propagates the probability density for a short time period by the system model (3), while the filtering step corrects the probabilities by the Bayes formula where the likelihood incorporates the measured data (4).

Now, we describe the recursive procedure. Suppose we have an estimate of a conditional probability *p*(*x*_t-1_|*y*_1:t-1_), where *y*_1:t-1_ stands for the series of measurement data *y*_1_, *y*_2_,,, *y*_*t*-1_. With this probability, we can estimate the probability *p*(*x*_t_|*y*1:_t-1_) at time *t*, using the short time propagation *p*(*x*_t_|*x*_t-1_) by the system model (1), as

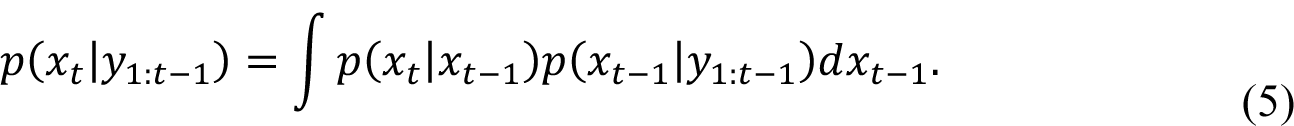

Here, we assumed the Markovian property of *x*_t_. This process is called the “prediction”. Then, with the predicted distribution *p*(*x*_t_ |*y*_1:t-1_), we can take into accounts the new measurement at time *t* as the likelihood *p*(*y*_*t*_ |*x*_*t*_), in the following formula,

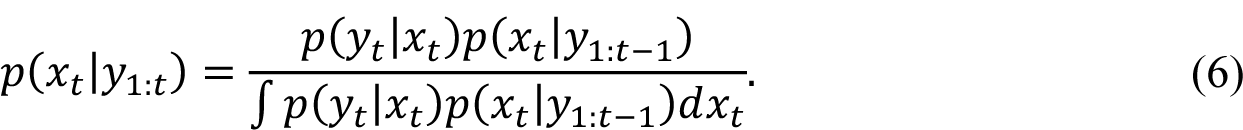

With (5) and (6) together, we can update the probability distribution by a unit time taking into a new measurement account. Thus, the iterative use of these equations results in the estimate of *x*_*t*_ given the experimental data *y*_1:*t*_.

For the LDS, we can calculate (5) and (6) analytically: Assuming the initial probability distribution is a normal distribution, we know all the probability distributions in (5) and (6) take Gaussian forms as well. Thus, we only need to obtain the formula for the update of the mean vector and the variance-covariance matrix to characterize these Gaussian distributions. For the prediction step (5), we can derive the updates in the mean vector and the variance-covariance matrix, respectively, as

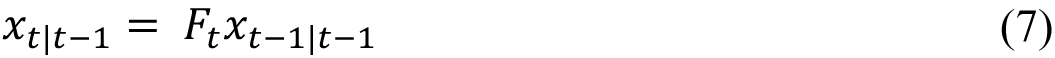

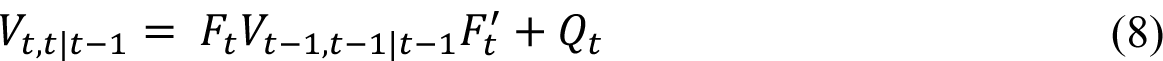

where the abbreviated subscript *t*|*t* − 1 for *x* and *t*, *t*|*t* − 1 for *V* mean that these are for *p*(*x*_*t*_ |*y*_1:*t*-1_) (Note that, for the matrix *V*, the notation of the doubled *t*, *t* is unnecessary/redundant in the Kalman filter method alone, but is necessary for the Kalman smoother method described later). We also note that the superscript’ for matrices represents the transpose of the matrix. For the filtering step (6), the updates in the mean vector and the variance-covariance matrix, respectively, can be written as

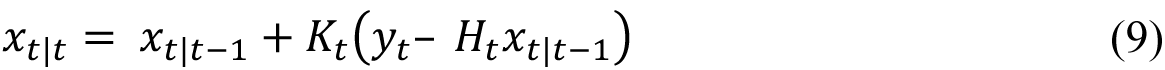

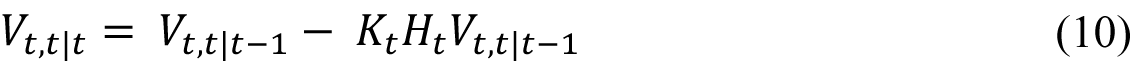

where the matrix, *K*_*t*_, called the Kalman gain, is defined as

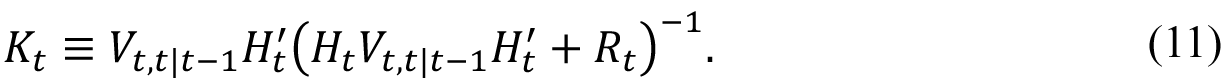

The iterative use of the prediction (7) and (8) and the filtering (9) and (10) results in the Bayesian inference of *x*_*t*_ given the experimental data *y*_1:*t*_ for the case of LDS, which is called the Kalman filter method.

### The Kalman smoother method

We note that the Kalman filter method uses the current and the past measurement data *y*_1:*t*_ to infer the probability of the current state *x*_*t*_, which is useful for the on-the-fly, i.e., real-time, inference. However, when measurement data for the whole process, *t* = 1, 2,,,*T* is available in the off-line mode, we may also use the “future” measurement data *y*_*t*1:*T*_, to infer the current state *x*_*t*_. The Kalman smoother method, or the RTS smoother (20), gives frameworks for this case. Among a few variants of the Kalman smoothers, we here focus on the fixed-point Kalman smoother (hereafter simply denoted as the Kalman smoother).

The Kalman smoother estimates the probability distribution of *x*_*s*_ at the fixed time point s for the measurement data *y*_1:*t*_ such that s < *t*. The Kalman smoother process starts with the normal Kalman filter method (7)-(10) up to obtaining *p*(*x*_*s*_|*y*_1:*s*_). To proceed further, we deal with *p*(*x*_*s*_, *x*_*t*_|*y*_1:*t*_), which becomes Gaussians distributions in the dual state space (*x*_*s*_, *x*_*t*_). For this dual state space, we can obtain recurrence formula similar to (7)-(10) for propagating *t*, in which the system model for the first variable *x*_*s*_ becomes the identity matrix (because the time s is fixed). Finally, by integrating out *x*_*t*_ (i.e., the marginalization), we obtain *p*(*x*_*s*_|*y*_1:*t*_).

Starting from *p*(*x*_*s*_, *x*_*s*_|*y*_1:*s*_) (equivalent to *p*(*x*_*s*_|*y*_1:*s*_)), in the prediction step for *t* (> the fixed time *s*), we can still use (7) and (8) to update the mean vector *x*_*t*_|_*t*-1_ and the variance-covariance matrix *V*_*t*_,_*t*_|_*t*-1_. The *x*_*s*_|_*t*-1_ needs not to be updated because of the identity matrix. In addition, we need to know cross-covariance matrix between the dual state spaces, i.e. *V*_*s*_,_*t*_|_*t*-1_, which is updated as

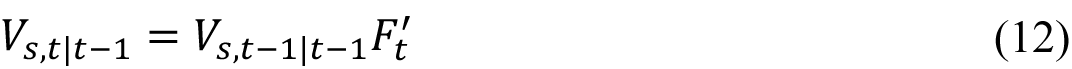

The filtering process needs, in addition to (9)-(11), extra formula to update the probability distribution of *x*_*s*_ by taking into accounts additional measurement *y*_*t*_,

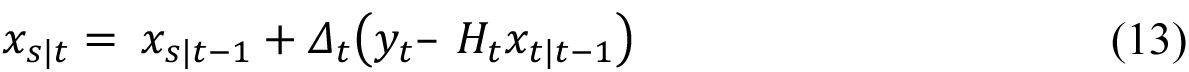

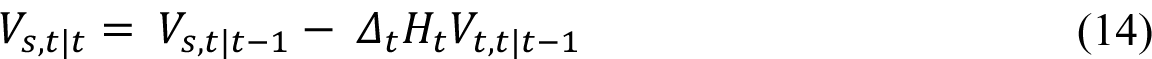

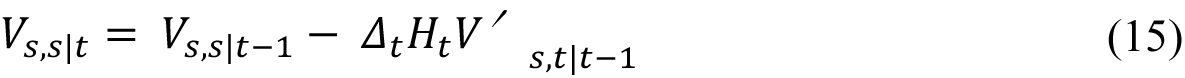

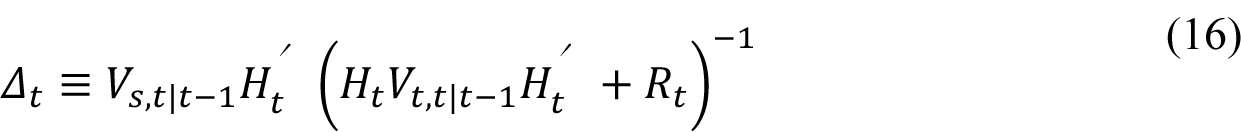

Altogether, we can iteratively proceed to obtain the mean vector and variance-covariance matrix for *p*(*x*_*s*_, *x*_*t*_|*y*_1:*t*_) for time *t* such that *t* > *s*.

### Protocol for HS-AFM

Now, we apply the Kalman filter and the Kalman smoother to the HS-AFM data, in which the height of target molecules on the stage surface, *xy*-plane, is measured via scanning the probe tip. The state vector *x*_*t*_ represents the height of target molecules at time *t* for all the pixel points *n*_*p*_ = *n*_*pX*_ × *n*_*pY*_. Thus, the state vector *x*_*t*_ has its dimension *k* = *n*_*p*_, where 2-dimensional image data are stacked into 1-dimensional vector. The HS-AFM measurement data at a time corresponds to the height of the molecule at one pixel. Thus, the measurement *y*_*t*_ is one dimension *l* = 1. (We note that, more in detail, the AFM data represent the height of the probe tip at which the finite-sized probe tip hits the target molecule, and thus is not identical to the height of the target. Since this difference does not alter the following argument in this study, we do not distinguish them for simplicity of description.)

For the system model (3), without prior knowledge on movements of target molecules, we use a minimal model. On the stage surface, biomolecules can be bound on the initial position with some Brownian motions. In the normal condition, the inertia can be ignored. Thus, as a default, we employ the minimal deterministic matrix *F*_*t*_ = *I*, the identity matrix. Later, we discuss the system model with non-zero diffusion. For the variance-covariance matrix of the system noise *Q*_*t*_, assuming that biomolecules have continuous and smooth shape, we introduce

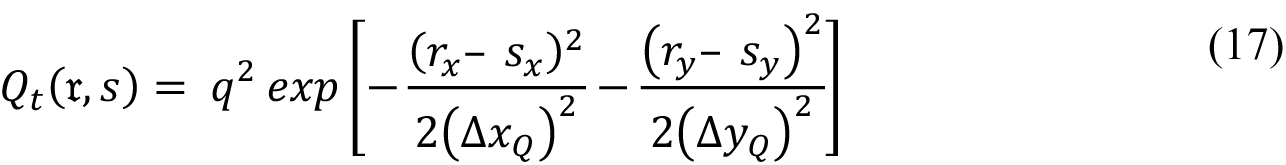

where *r* ≡ (*r*_*x*_, *r*_*y*_), *s* ≡ (*s*_*x*_, *s*_*y*_) and Δ*x*_*Q*_ = Δ*y*_*Q*_ = 1 pixel in this study for simplicity. The parameter *q* sets the scale of the spatial correlation.

The HS-AFM experiment measures the molecular height at a pixel per unit time. Thus, the measurement *y*_*t*_ is one dimension *l* = 1, and *H*_*t*_ in (4) is a 1 × *k* matrix, in which only one element that corresponds to a measured pixel at the time is unity and all the others zero. In this study, we assume the raster scan as the measurement schedule (Figure 1A): The probe scans in the *x*-direction always from 0 to n_px_-1. After one-line scan, the probe moves in the *y*-direction by one pixel and repeats the scanning in the *x*-direction. This is repeated until completion of the one-frame measurement. The variance-covariance matrix R_*t*_ was assumed to be diagonal, R_*t*_ = *rI* for simplicity. As a default, we set *r* = 1.

We illustrate the scheme using a simple case of an image with 3 x 3 pixels (Figure 2B). During *t* = 1, 2,,, 9, for example, the probe scans pixel by pixel for one frame image, measuring the specimen height. The raw AFM frame image obtained by the time *t* = 9 is defined as the collection of the scanned data over *t* = 1 ~ 9. Thus, the raw AFM images contain asynchronous measurement. For *t* = 1 ~ 9, the Kalman filter alternatively uses the prediction (7) (8) and the filtering (9)-(11) that uses the same data as the raw AFM case. While the Kalman filter can produce the whole image (the mean vector *x*_*t*_|_*t*_ in (9)) at every time *t* = 1, 2,,, 9, we use only *t* = 9 point, *x*_9|9_, for comparison with the raw AFM image. For the same time point s = 9 as the fixed-point, we design the Kalman smoother with use of the data for *t* = 1 ~ 18, obtaining *x*_9|18_ in (13). Notably, the Kalman filter estimates the image *x*_*t*_ using the data at the same and past time, whereas the Kalman smoother in the current form estimates the image *x*_*t*_ using the data for the upcoming one scan as well as those in the past.

In the twin-experiment for the toy model, we compare raw AFM movies, the movies from the Kalman filter and the Kalman smoother with the ground-truth movie. The latter is defined as the molecular height exactly at the right time. In the illustration of 3 x 3 pixels, the ground-truth frame image corresponds to the specimen height for all the pixels at *t* = 9. Note that, in real AFM experiments, we cannot obtain this ground-truth movie.

In the twin-experiment for the conformational change of dynein, we use the image size of 25 x 40 pixel in *xy*-directions. From a MD trajectory reported previously (23), we prepared 25,000 snapshot structures. With use of snapshots at every 1000 time points, the ground-truth AFM-like frame images are produced by a software afmize (24). Thus, the ground-truth movie contains 25 frames. We mimic the HS-AFM measurement by obtaining the molecular height data at one pixel per MD snapshot. Thus, after 25 x 40 = 1000 observations, we obtain one complete frame image that mimics the HS-AFM. We obtain 25 frames of HS-AFM movie mimic. While the Kalman filter method can produce the AFM-like images at every time points, we produced the corresponding 25 frames with the interval of 1000 time points. The Kalman smoother method in the current setup uses 1000 extra-observations in future as well as the current and past data, and thus the image cannot be generated for the final 1000 time points. Using the same time interval of 1000 time points, we thus generated 24 frames by the Kalman smoother. For the method comparison, we used the 24 frames with the interval of 1000 time points.

### The correlation coefficient

In the examination, to quantify the resemblance of the reconstituted AFM image with the ground-truth AFM-like image, we used the modified correlation coefficient(c.c.) used in earlier works (24–26). The c.c is defined as

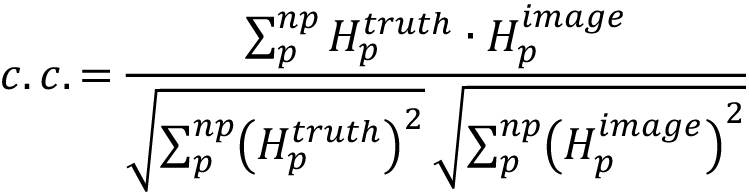

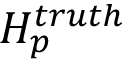 and 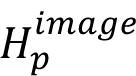 are, respectively the ground-truth and reconstituted heights at the pixel point, p.

## Results

### A toy cone model

We begin with a toy model made of a randomly diffusing cone shape (Figure 3A, Movie S1). We set the observation area of 10×10 pixels. The cone has its base radius 3 pixels and the height 3 (the height is in an arbitrary unit). The position of the cone vertex makes a random walk for 10000 steps within the observation area. In each discrete time interval, the cone position moves independently in *x* and *y* directions by the moves randomly drawn from the uniform distribution in the range *s* × [−1, 1), where *s* is a scaling factor set as either 1, 0.1, or 0.01, for fast, default, or slow diffusion of the cone. At the edges of the observation area, the move is bound to inside.

**Figure 3.**
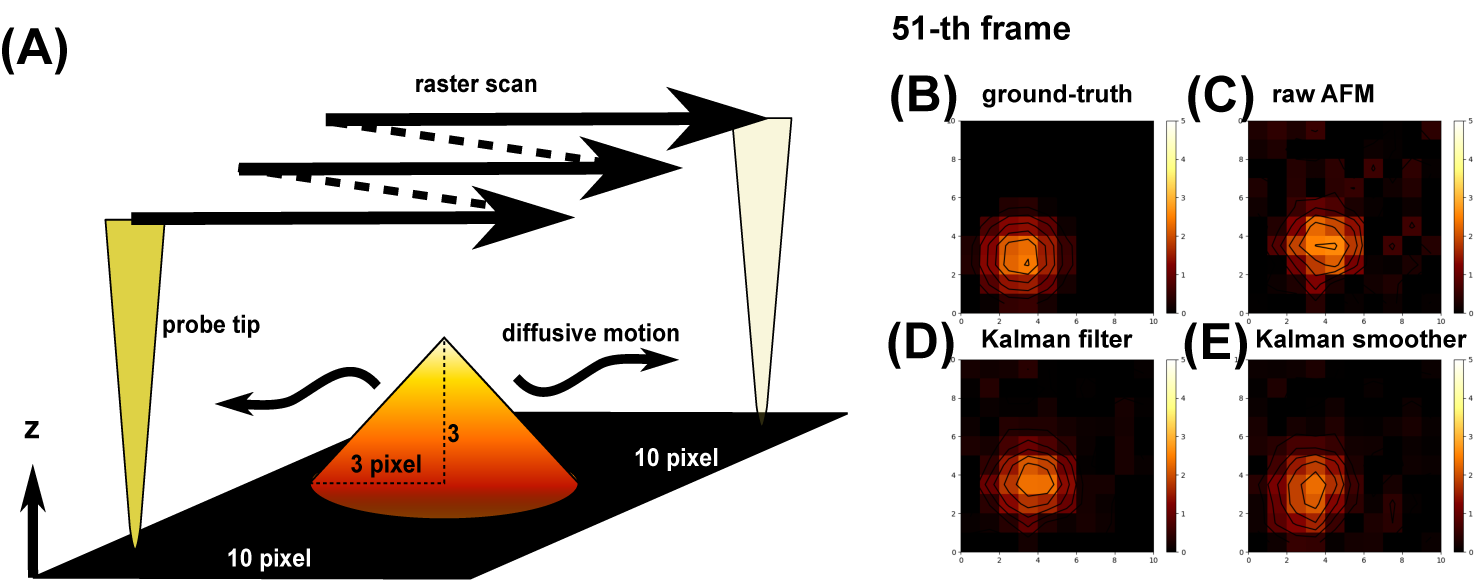
A toy diffusing cone model. (A) Cone shape and stage setup. AFM-like images at 51-st frame are shown for (B) the ground-truth data, (C) the mimic of raw AFM image, (D) the Kalman filter image, and (E) the fixed-point Kalman smoother image. (B-E) Color maps are shown on the right side (matplotlib, afmhot). The black curve represents the contour plot.

The HS-AFM measurement is mimicked to get the height data at one pixel per discrete time by the raster scan (we use the height of the cone at a representative position of each pixel). Thus, it takes 100 discrete times to obtain one frame image of 10 x 10 pixels. The 10000-step random walk results in an AFM-like movie that contains the 100 frames. We added a spatiotemporally uncorrelated Gaussian noise with the mean zero and the standard deviation 0.3 to the height data.

Using the diffusing cone with the default diffusion coefficient (*s* = 0.1), we created the raw AFM movie (Movie S2), the Kalman filter movie (Movie S3), and the Kalman smoother movie (Movie S4), as well as the ground-truth movie (Movie S1). The ground-truth movie is defined as the series of images that record the cone heights in all the pixels at once (Note that this is possible only in twin-experiments, but not for real HS-AFM experimental data). Figure 3B, C, D, and E show the snapshots for the 51-st frame image in the ground-truth, the raw AFM, the Kalman filter (the system noise *q* = 0.1), and the Kalman smoother (*q* = 0.1), respectively. At first sight, we see that the raw AFM image is apparently noisy and the image near the vertex is clearly distorted. In comparison, the Kalman filter and smoother give less noisy images with less distorted shapes around the vertex. Therefore, both the Kalman filter and smoother method seem to reduce the noise and the data asynchronicity, which we will quantify below.

To quantify the similarity to the ground-truth movie, we calculated the average correlation coefficient (c.c.) over 99 frame images for the raw AFM movie, the Kalman filter movie, and the Kalman smoother movie with different values of the parameter *q* (Figure 4A, data sorted by the value of c.c.)(Note that the final time *t* = 100 is discarded because the Kalman smoother does not produce the image at the final time point).

**Figure 4.**
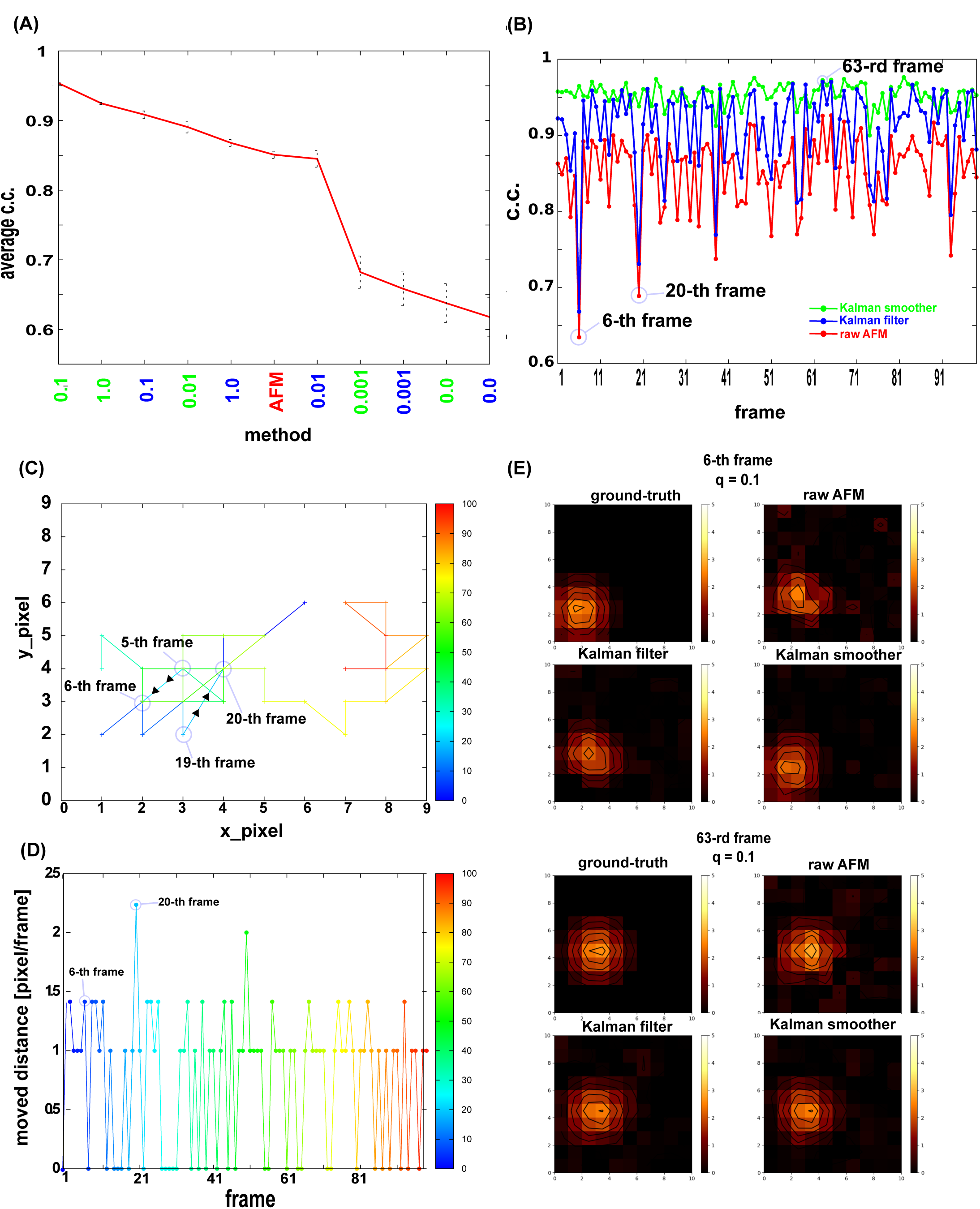
Quantitative comparison of the Kalman filter and smoother methods with the raw AFM image for the toy cone model. (A) The correlation coefficients (c.c.’s) with the ground-truth movie averaged over time for the Kalman filter (blue) and the Kalman smoother (green) results with different q-values, together with the case of the raw AFM movies (sorted by the c.c. value). The error bar represents the standard deviation of the average estimate. (B) Time series of c.c.’s for the raw AFM movie (red), the Kalman filter with q = 0.1 (blue), and the Kalman smoother with q = 0.1 (green). In the raw AFM movie, the lowest c.c. was found at the 6-th frame. (C) The trajectory of the pixel position at which the cone peak is located. The color represents the frame; blue for the initial and red for the final frames. (D) The travel distance between the two frames, in the unit of edge length of each pixel. (E) Comparisons of AFM-like frame images at which the average c.c.’s were lower and higher in the raw AFM movie, the 6-th and 63-rd frames, respectively. For each, results of the ground-truth (top left), the raw AFM (top right), the Kalman filter with q = 0.1 (bottom left), and the Kalman smoother with q = 0.1 (bottom right) are shown.

In this example, the largest c.c., 0.95, is achieved by the Kalman smoother with *q* = 0.1. The best of the Kalman filter was realized with *q* = 0.1 too, of which c.c. value is ~0.91. The raw AFM movie has the c.c. ~0.85, markedly lower than the best Kalman smoother movie. We note that the standard deviation of the c.c. tends to increase as the average c.c. decreases, implying that methods with lower average c.c.’s may fluctuate more along time.

Therefore, next we plot the time series of c.c. of the Kalman filter, the Kalman smoother with the best parameter *q* = 0.1, as well as the raw AFM data in Figure 4B. Among the three curves, we see that the Kalman smoother method clearly outperforms with nearly always and stably the best c.c.’s. The Kalman filter shows some improvement over the raw AFM data. Interestingly, the raw AFM data exhibit the largest fluctuation along time; it sharply drops occasionally, such as the 6-th frame and the 20-th frame. The Kalman filter showed the same tendency as the raw AFM data.

This instability can be understood by looking into the trajectory and the moves of the pixel at which the cone vertex exists (Figure 4CD). Figure 4C shows that the cone made a large step from the 19-th to the 20-th step. From the 5-th to 6-th, the move distance was not necessarily the largest (Figure 4D), but the cone moved to the right, which is opposite to the scanning direction. At these instances, the probe tip was not precisely observing that pixel, resulting in larger deviation. The raw AFM image at the 6-th frame is clearly distorted with a shifted vertex position (Figure 4E, top right). At the same frame, the Kalman filter image is less noisy, but the vertex position is significantly shifted to the top-right direction (Figure 4E bottom left, the peak at the pixel (2,3)), which corresponds to the position at the previous, i.e., 5-th, frame. Notably, the Kalman smoother did not show marked drops in c.c. in these frames (Figure 4B) and the image at the time looks much closer to the ground-truth image (Figure 4E bottom right).

In contrast, at some frames, the raw AFM, the Kalman filter, and the Kalman smoother all show accurate images with high c.c.’s, such as the 63-rd frame (Figure 4B). The images shown in Figure 6E suggest that all the images are rather similar. This is primarily because the cone did not move much from the 62-nd to the 63-rd frames.

So far, we showed that the Kalman filter and smoother methods reduce the image distortion in the raw AFM image, which can contain the two aspects; the resolution of the data asynchronicity and the noise reduction. To better distinguish the two aspects, we apply the Kalman filter and smoother to the noiseless version of the diffusing cone model.

We compare the performance for this noiseless AFM-like movie in Figure 5, where we find that the Kalman smoother outperforms the Kalman filter and the raw AFM movie in the same way the previous case with the noise (Figure 5A). Comparing the result by the Kalman filter and the raw AFM movie, we see nearly equivalent level of average c.c.’s (Figure 5A), as well as the time series of c.c. (Figure 5B). This suggests that the major effect of the Kalman filter in the previous case with the noise is the noise reduction, but not the resolution of the data asynchronicity. In contrast, the Kalman smoother steadily improves the image quality in the time series data (Figure 5B). Looking the images in Figure 5C, we find that, upon large-scale motions of the cone, the Kalman smoother can catch up the motion with the vertex position about the same, showing its performance in the resolution of the data asynchronicity. On the other hand, the raw AFM and the Kalman filter images tend to have their peaks in the old positions.

**Figure 5.**
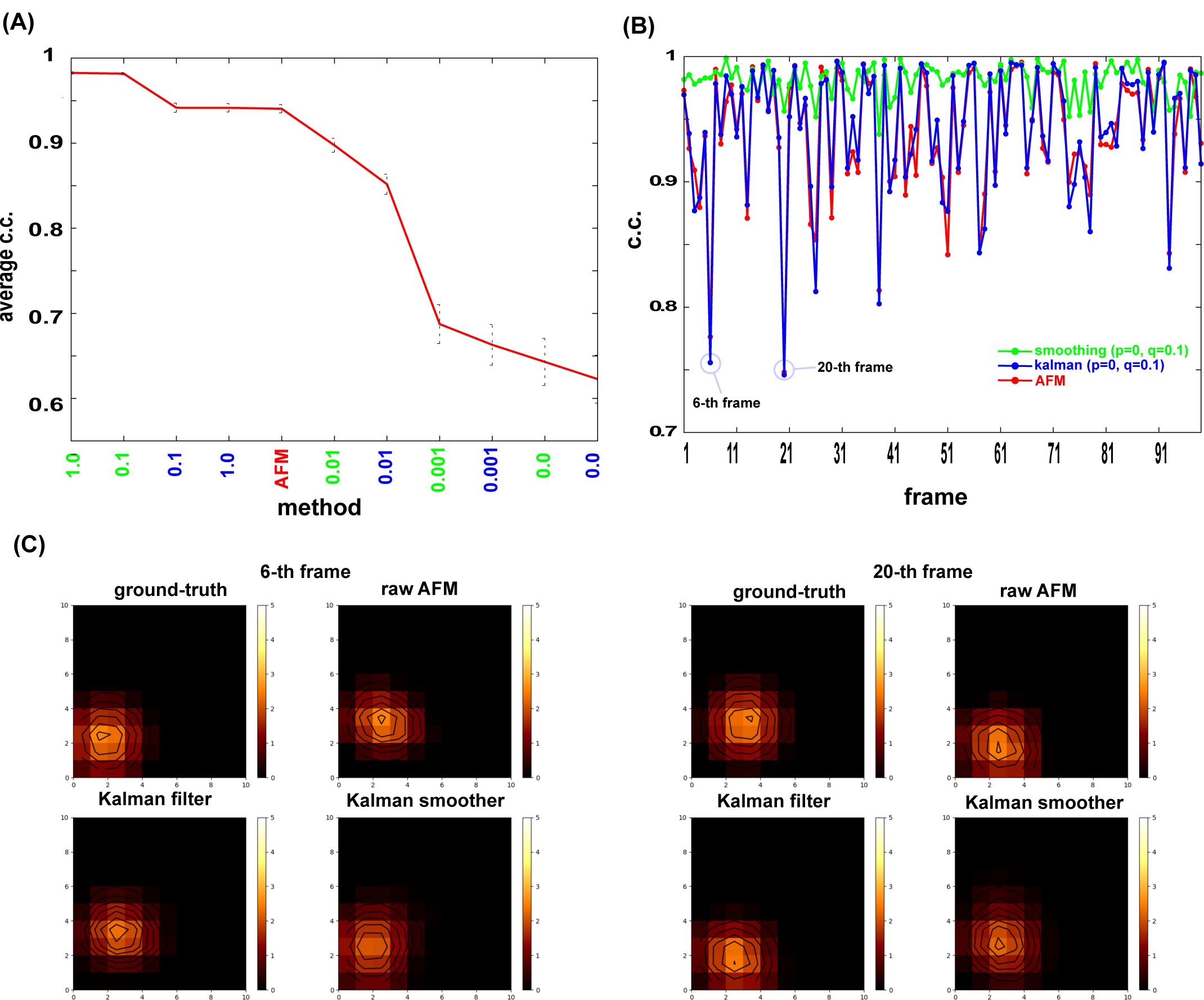
Quantitative comparison of the Kalman filter and smoother methods with the raw AFM image for the toy cone model without Gaussian noise. (A) The same as Figure 4A. (B) The same as Figure 4B. (C) The same as Figure 4E.

So far, we used the diffusing cone model with the default (medium) diffusion coefficient (scaling factor s = 0.1). What happens for faster and slower diffusion, relative to the probe scanning time. With use of the scaling s = 0.01 for the slow diffusion and s = 1 for the fast diffusion, we repeated otherwise the same analysis (with the noise). For the slowly diffusing cone movie, we see the Kalman filter works nearly as much as the Kalman smoother (Figure 6). This is because the data asynchronicity does not become serious issue in such a slowly moving specimen. Noticeably, a use of smaller system noise q = 0.01 was slightly better than q = 0.1 which was the optimal in the case of the default diffusion.

**Figure 6.**
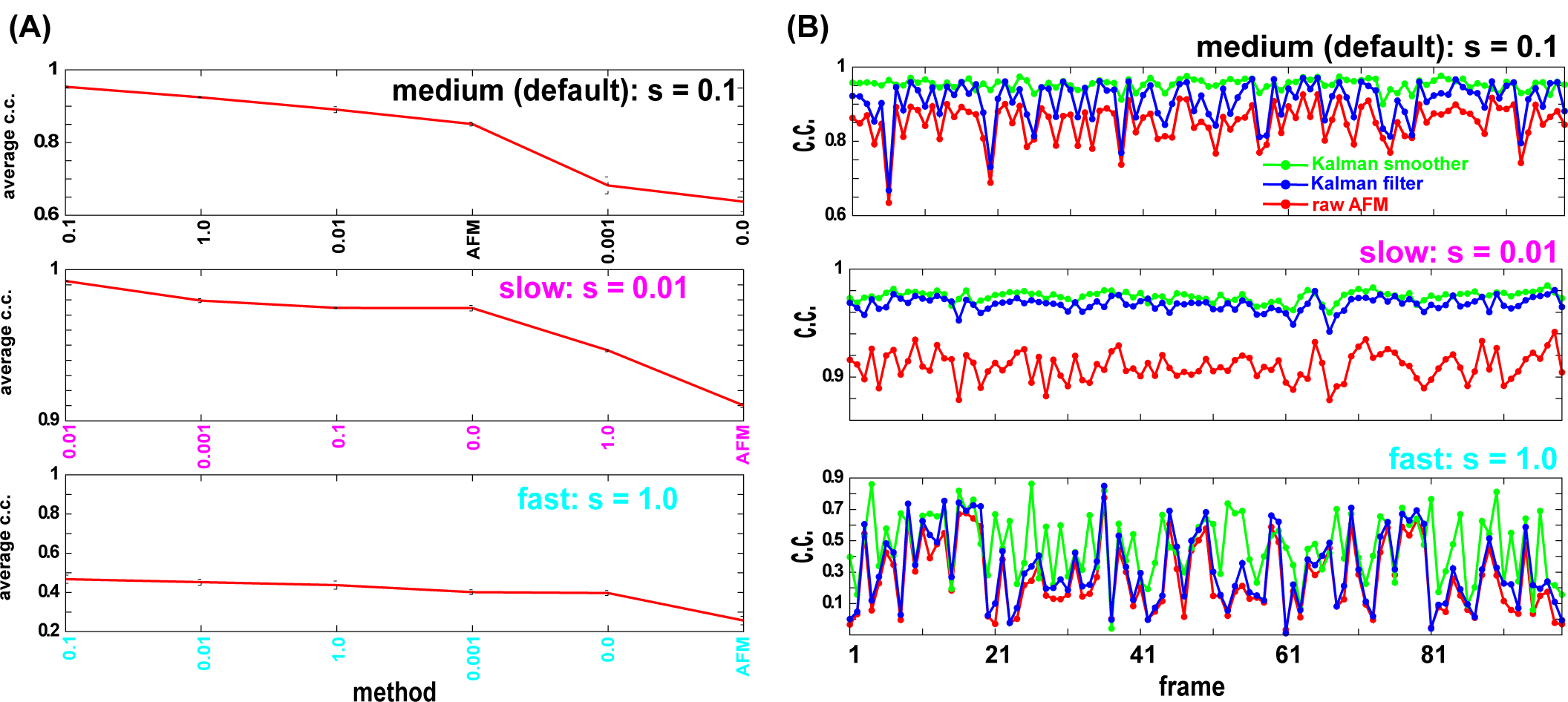
Quantitative comparison of the Kalman filter and smoother methods with the raw AFM image for the toy cone model with different diffusion rate. (A) The c.c.’s with the ground-truth movie averaged over time for the Kalman smoother results with different q-values, together with the case of the raw AFM movies (sorted by the c.c. value). The results for the medium (default) diffusion rate (s = 0.1, top), the slow diffusion rate (s = 0.01, middle), and the fast diffusion rate (s = 1, bottom) are shown. (B) The same as Figure 4B for the medium, slow, and fast diffusion rates. Both the Kalman filter and smoother use q = 0.1.

For the fast diffusion case, the Kalman smoother, the Kalman filter, and the raw AFM movies all produced poor movies. Since the data asynchronicity is the major issue, the performances of the raw AFM and the Kalman filter are nearly equal. The Kalman smoother showed the highest c.c. among the three cases, albeit with rather low c.c.’s in many time points (Figure 6B).

### A twin experiment for protein conformational change; dynein

Next, we examine a conformational change of a motor protein, dynein, with a twin experiment. Namely, using the molecular model and a molecular dynamics simulation tool, we previously performed molecular dynamics simulation of relatively large-scale conformational changes of dynein (23). From one trajectory obtained previously, we made the ground-truth movie, the synthetic AFM movie that mimics the HS-AFM measurement with data asynchronicity, and the movie via the Kalman smoother using the same HS-AFM measurement data.

Cytoplasmic dynein, dynein for brevity, is an ATP-dependent motor protein which forms a homo-dimer and “walks” on the microtubule filament serving as a transporting machine in cells. (Figure 7A left). Coupled with the ATP hydrolysis cycle, the motor domain of dynein makes characteristic conformational change (Figure 7A right). Among many changes, the most prominent conformational change is found at the linker which connects the motor domain with the cargo and the other monomer (drawn in purple in Figure 7A left and right). In this conformational change, called the power-stroke, the linker swings from a highly bent form (in the pre-state) to a straight form (in the post-state). This swing is supposed to be responsible for the motility of dynein. In addition, the stalk, which is a long extrusion from the ring structure (grey in Figure 7A right) also changes their shape. Previously, we performed molecular dynamics simulations of the power-stroke of the motor domain of dynein (23), one of which is used for the analysis here.

**Figure 7.**
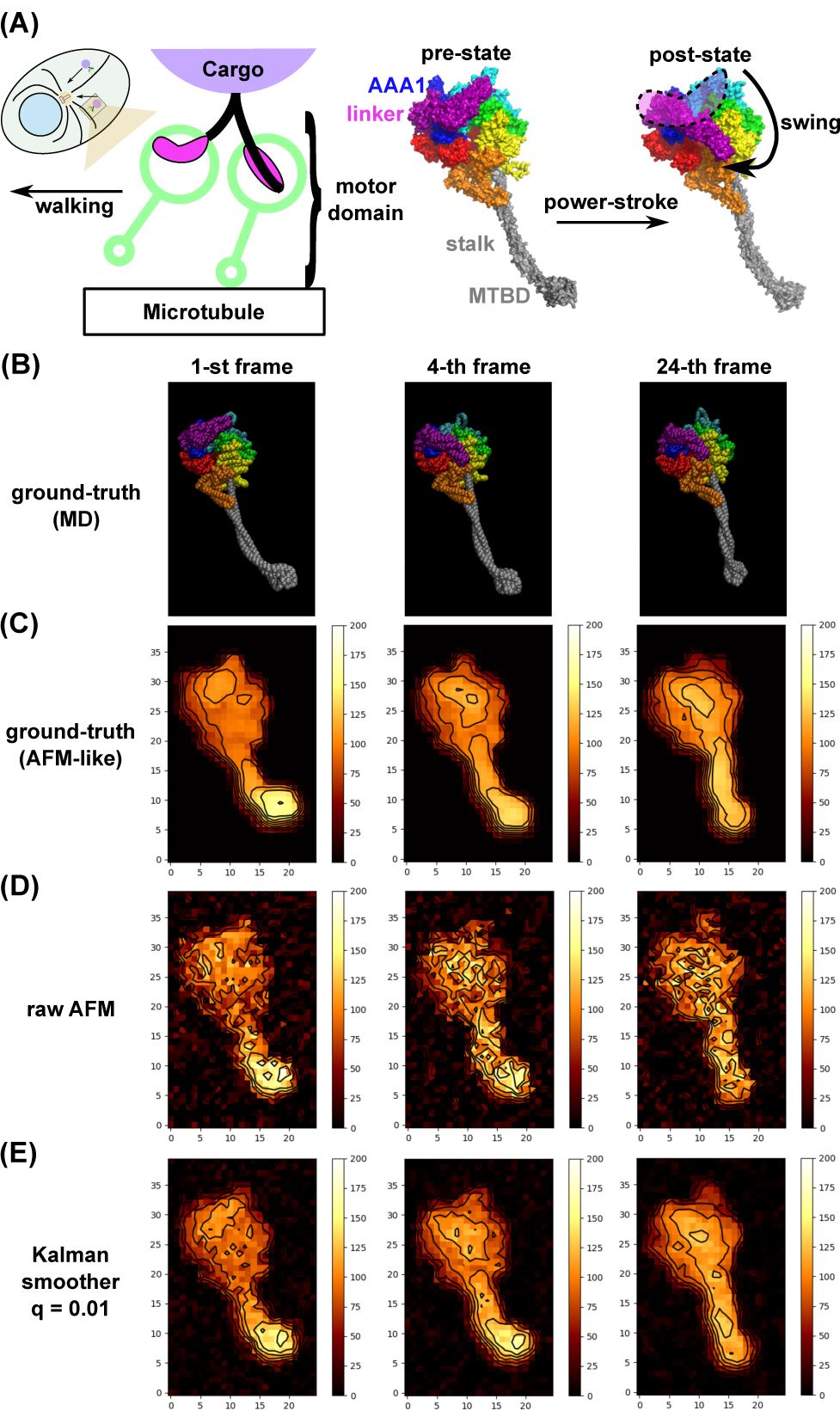
Kalman smoother for a synthetic AFM movie of a motor protein, dynein. (A) Structure of dynein. (left) The dynein is a homo-dimeric motor protein that transports cargos along microtubule driven by ATP hydrolysis. (right) During the ATP hydrolysis cycle, dynein exhibits the power-stroke motion where the linker (purple) changes from a bent form (the pre-state) to a straight form (the post-state), in addition to the structure change in the stalk/MTBD, the extruded region (grey). (B-E) The dynein structure and the corresponding AFM-like images are depicted for the initial state (the 1-st frame, before the power-stroke), for the 4-th frame, after the power-stroke), and at the end (the 24-th frame). (B) Snapshot structures from a MD trajectory. (C) The ground-truth image produced from MD snapshots. (D) The mimic of raw HS-AFM images. The Gaussian noise is added to the data obtained one pixel per measurement. (E) The Kalman smoother (q = 0.01) applied to the data in (C). The stage corresponds to the zero height (black). Contours are added to emphasize the differences among methods.

Compared to the previous toy model of diffusing cone, our focus here is whether we can observe the conformational change of dynein molecule, especially the linker power-stroke motion, via the Kalman smoother. Since such a conformational change is relatively fast, the data asynchronicity in HS-AFM measurement must be a more serious issue.

The MD trajectory starts from the pre-state (the 1-st frame in Figure 7B), and shows the power-stroke from the first to the third frames, which is followed by the conformational change in the stalk (as depicted in the last (24-th) frame). Notably, the power-stroke is a rapid motion and occurs between the second and third frames. Using these snapshot structures, we made the series of noiseless instantaneous AFM-like images, the ground-truth movie (Figure 7C Movie S5) (25 x 40 pixels, 1 nm per 1 pixel); we used the standard collision-detection algorithm with the tool, afmize (24), to estimate the molecular height for all the pixels at once. With the ground-truth movie, we can easily recognize the power-stroke, as well as the stalk structure change.

Next, we created a movie that mimics the HS-AFM measurement. We first added the spatiotemporally uncorrelated Gaussian noise with the mean zero and the standard deviation 2nm in the height, and made the movie that mimic HS-AFM measurement; recording the height of one pixel per measurement with the raster-scanning (Figure 7D Movie S6). Due to relatively large noise and rapid conformational change, the power-stroke motion is hardly identified.

Then, we applied the fixed-point Kalman smoother to the same measurement data, with q = 0.01 (Figure 7E, Movie S7). Apparently, the resulting movie clearly exhibits the power-stroke motion at the 4-th frame and the stalk conformational change by the last frame, albeit some distorted images.

Varying the parameter q in the system model, we quantitatively compared the results of the Kalman smoother, in addition to the raw HS-AFM data, using the average correlation coefficients (c.c.). The result in Figure 8A shows that the Kalman smoother with q = 0.01 gives the best average c.c. value (~0.978), which is significantly higher than the result of raw AFM case (~0.914).

**Figure 8.**
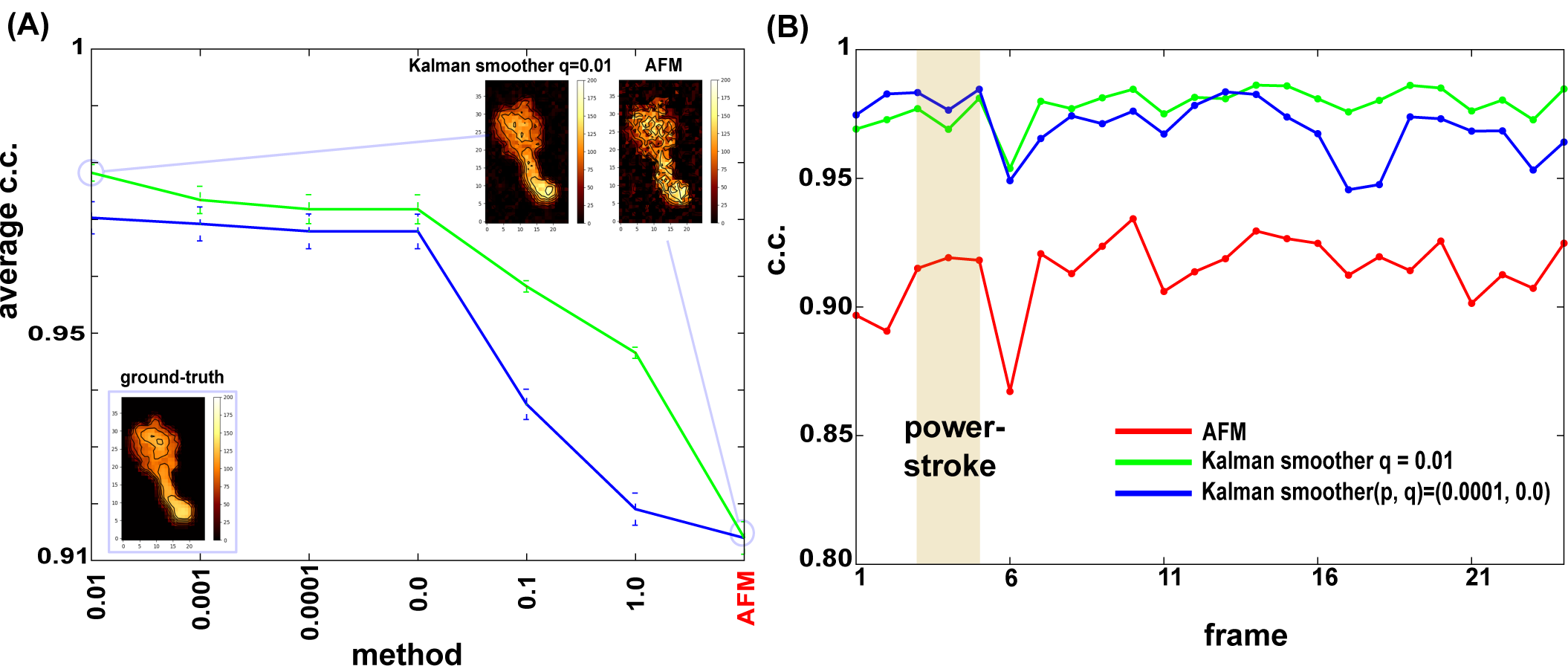
Quantitative comparison of the Kalman smoothers with different parameters. (A) The mean of the correlation coefficient (c.c.) of the fixed-point Kalman smoother with different *q* parameters (the diffusion parameter), in addition to the raw synthetic HS-AFM movie, for dynein. The data are sorted in the descending order of the mean c.c. The bar represents the standard error. (B) The time series of c.c. In (A) and (B), red; the raw AFM movie, green; (p=0, q=0.01), blue; (p=0.0001, q=0.0). In (B), the brown shaded box; the period of the power-stroke.

Figure 8B shows time series of c.c. for the raw HS-AFM images and Kalman smoother images. We clearly see that, throughout the entire process, the Kalman smoother improves the image quality significantly. Notably, the c.c. of the raw HS-AFM fluctuates largely, while that of the Kalman smoother is more stable. All the c.c.’s drop at the 6-th frame, which is due to a rapid conformational change in the stalk/MTBD region, as will be discussed below.

Here, we tested the case with non-zero diffusion in the matrix *F*_*t*_ of the system model. We model the diffusion by introducing a small probability, *p*, in off-diagonal elements that correspond to the shift in one-pixel to either left, right, up, and down directions. A large value of *p* washes out all the feature immediately. A small non-zero value of *p*, sometimes improve the produced image. Here, we show the results with p = 0.001 together with *q* = 0. While the use of non-zero p values does not increase the average c.c. compared to the p = 0 case (Figure 8A), the time series of c.c. suggest that a non-zero p value can improve c.c. at early time range (Figure 8B). Non-zero *p* value tends to spread the probability distribution, in general. In the case of highly noisy raw image, this spread has the denoising effect, thus improving the quality of images. This denoising effect could be slightly higher than a similar denoising effect from non-zero *q* value. However, as time goes, the non-zero *p* value tends to spread the signal itself, reducing the c.c. in later stage.

Next, focusing on the power-stroke motions around the linker as well as the stalk/MTBD change, we compare the ground-truth movie, the raw AFM mimic movie, and the Kalman smoother movie more closely (Figure 9, the second line gives the close-up views of the linker). In the ground-truth movie, the 2-nd frame clearly indicated the pre-power-stroke configuration, whereas the 3-rd frame is in the middle of conformational change (Figure 9A). The 4-th frame corresponds to the post-power-stroke configuration. The ground-truth movie also shows that, from the 5-th to the 6-th images, stalk/MTBD change their conformation. Apparently looking at the raw AFM movie, we do not notice such a subtle conformational change at all (Figure 9B). In contrast, the movie produced from the Kalman smoother clearly reproduces the feature of power-stroke, albeit rather noisy images (Figure 9C). The 2-nd frame captures the pre-power-stroke configuration. The 3-rd and 4-th frames look like post-power-stroke. Not only the identification of the power-stroke but also its timing is correctly reproduced by the Kalman smoother method. As for the stalk/MTBD change, while the Kalman smoother identify the change between the 5-th frame and 7-th frame, its image at the 6-th frame resembles to the pre-structure change, which clearly deviates from the ground-truth movie (Figure 9D depicts the deviation; red (blue) where the Kalman smoother image has smaller (larger) height). Thus, the structure change in the stalk/MTBD was too fast to be detected by the Kalman smoother method. Interestingly, we also note that the Kalman filter method produced the movie that contains detectable power-stroke motion, but its time is retarded from that in the ground-truth and the Kalman smoother (data not shown).

**Figure 9.**
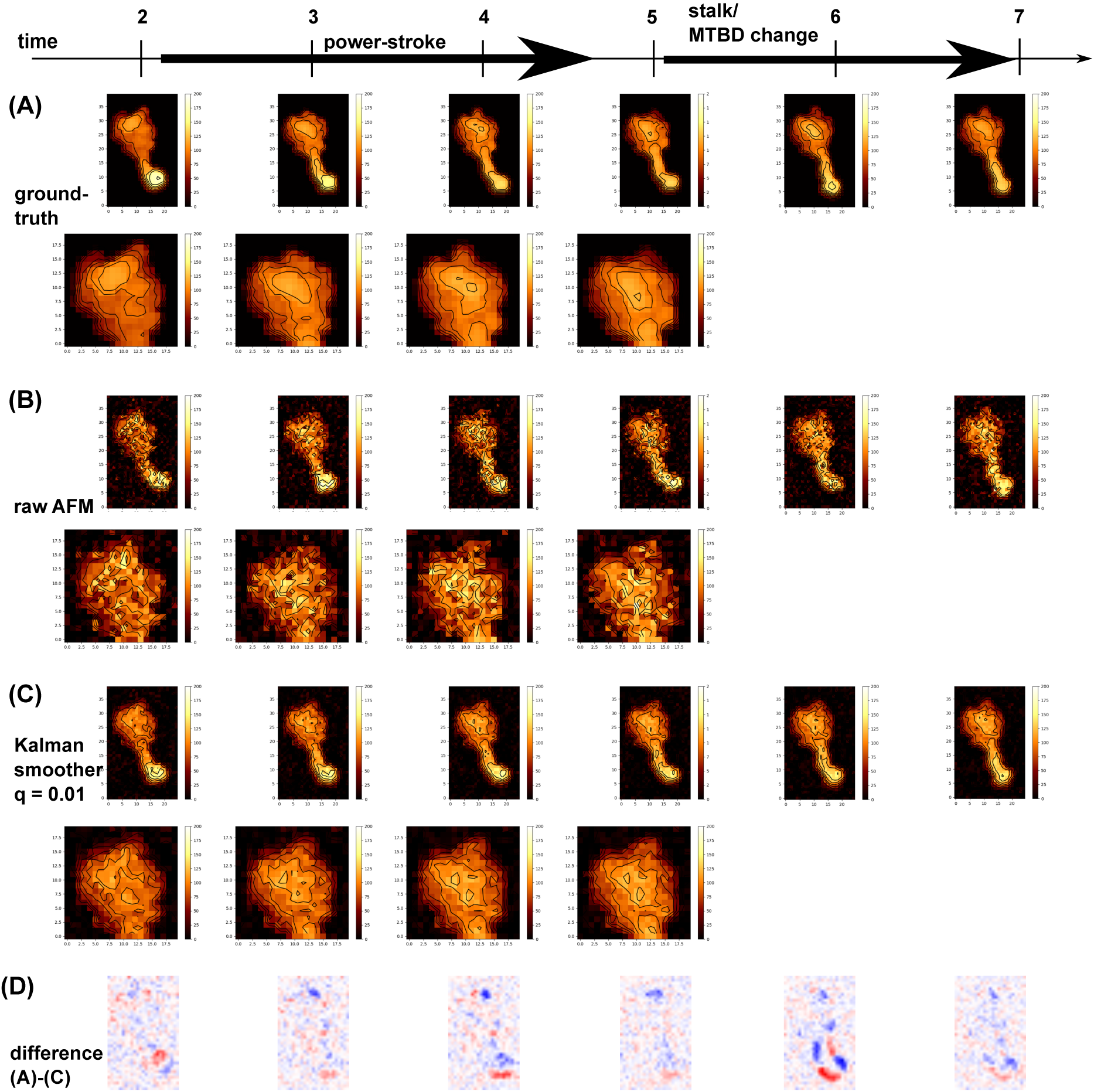
Comparison of AFM-like images in the power-stroke and stalk/MTBD change in dynein. The 2-nd to 7-th frame images of the ground-truth from a MD trajectory (A), the mimic of the raw AFM (B), and the Kalman smoother with q = 0.01 (C). In each case, the first line is for the entire image, while the second line magnifies the region where the linker swings during the power-stroke. The power-stroke occurs from the 2-nd to the 4-th frames. The stalk/MTBD change occurs from the 5-th to the 7-th frame. The stage corresponds to the zero height (black). Contours are added to emphasize the differences among methods. (D) The difference image between the ground-truth (A) and the Kalman smoother (C) images. Red and blue represent that the Kalman image has smaller and larger heights that the ground-truth image, respectively.

### Kalman smoother for real HS-AFM movie; a flagellar biosynthesis protein

Now, we apply the Kalman smoother method to real HS-AFM movies. The first example is the cytoplasmic domain of a flagellar biosynthesis protein, FlhAc, for which recent HS-AFM measurements revealed structural dynamics (27). Many bacteria, such as *Escherichia coli* and *Salmonella,* swim in fluid environment using a long filament called flagellum (Figure 10A left top). In its biosynthesis, many flagellar proteins are transported from the inner cell to the base of flagellum, and are piled up outside the cell membrane sequentially. This biosynthesis system includes a transporting machinery together with a gate. The target protein, FlhA, makes a ring form with nine copies of the molecules, serving as the gate of transport (Figure 10A center). The high-resolution structure of the cytoplasmic domain of FlhA, denoted as FlhAc, has been solved previously (28). It is composed of four domains (blue, green, yellow, and red in Figure 10A right bottom), in which each of two domains (blue/green, and yellow/red) forms a tight globular lobe, and two globular lobes are loosely connected via a flexible linker (depicted as two ovals and a line in Figure 10A right bottom).

**Figure 10.**
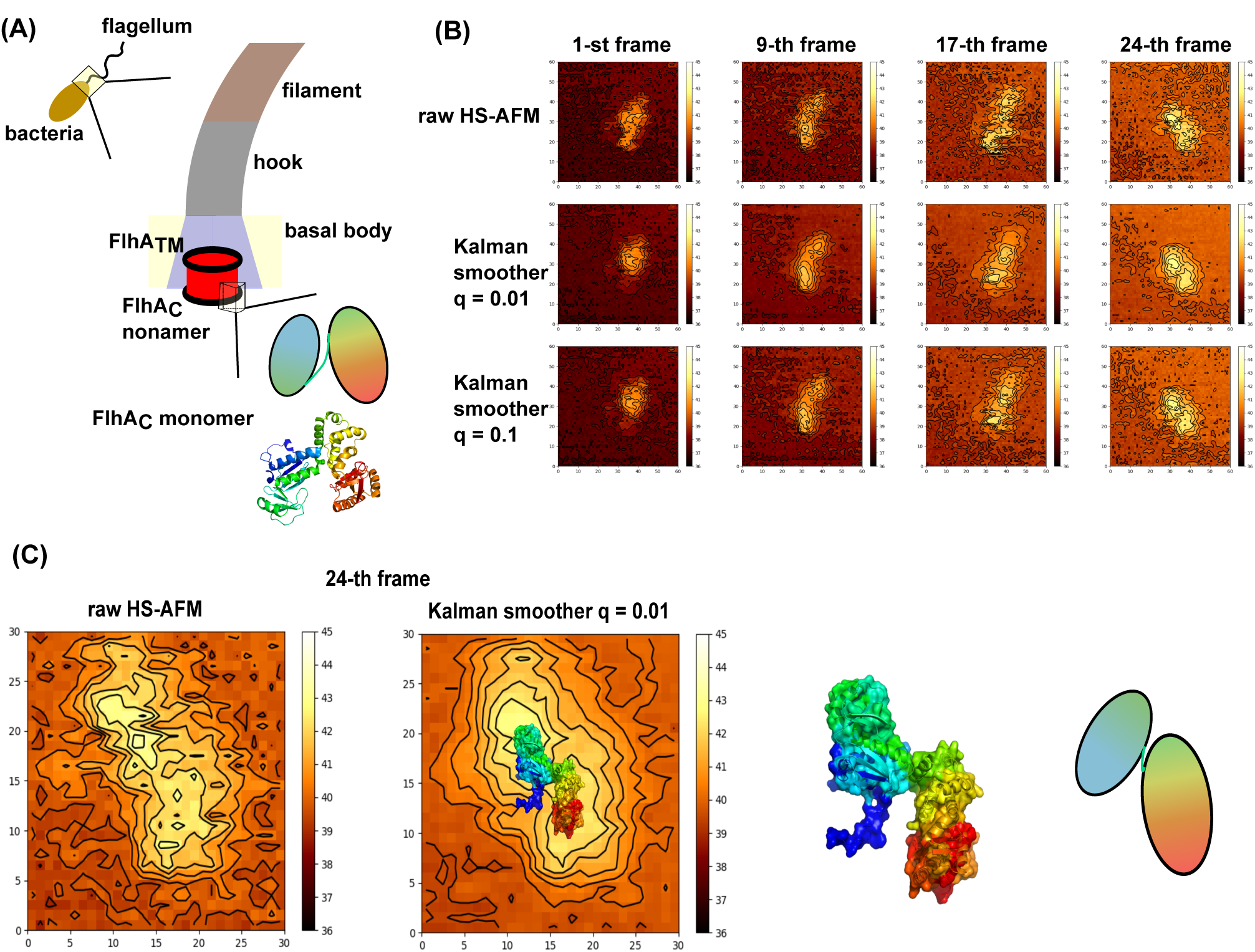
The Kalman smoother for a real HS-AFM movie of FlhA_C_. (A) Many bacteria swims using a tail-like flagellum (left). Proteins that constitute flagellum parts, filament, hook, and basal body, are transported from the inner cell via a gate which is made of nonamer of FlhA (middle). The monomeric cytoplasmic domain of FlhA, FlhAc, contains two lobes connected via a flexible linker, and its high-resolution structure is available (right). (B) Some frames (1-st, 9-th, 17-th, and 24-th) of raw HS-AFM movie (top), movies obtained by the Kalman smoother with q = 0.01 (middle), and q = 0.1 (bottom) are shown. (C) For the 24-th frame, a modeled conformation of FlhAc is superimposed on the image by the Kalman smoother (the second from the left). The conformation was obtained by rotating the flexible linker between two lobes (the right two pictures).

Structural dynamics of FlhAc was recently observed by the HS-AFM (27) (some frames depicted in Figure 10B top row)(Movie S8), in which we can guess the presence of two globular lobes, albeit significant noise. In addition, molecular overall shape looks somewhat distorted in some frames. We applied the Kalman smoother with two different parameters q = 0.01 (Figure. 10B, the middle row, Movie S9), and q = 0.1 (the bottom row) to a short fragment of a movie containing 26 frames (denoted as the 0-th to 25-th frames) of 60×60 pixels. The results in Figure 10B clearly show that the Kalman smoother markedly reduces the noise and identifies the two globular shapes more clearly. Oval shapes of individual globular parts are less distorted. Especially, we find visually that the parameter q = 0.01 works best in this case.

Encouraged by the clearer envelops in the Kalman smoother movies, we try to align a molecular structure of FlhAc to an image (Figure 10C). We assumed that the edge length of 1 pixel corresponds to 0.983 nm. Rigidly rotating the previously obtained structure model of FlhAc (the protein data bank id = 3A5I chain A), we repeatedly calculated the AFM-like images of the model structure, using afmize with the probe tip radius 1 nm and the apex angle 10º. We, however, could not obtain a reasonable fit. As described above, FlhAc is composed of two globular shapes connected via a flexible linker. We thus changed the conformation of the linker i.e., via rotation around phi/psi angles, seeking a better fit. The resulting FlhAc structure and the AFM-like image from this structure is shown in Figure 10C. Note that the alignment here is merely by visual inspection, and thus not very quantitative. More quantitative fit would require the flexible fitting as was recently reported elsewhere (24).

### Kalman smoother for real AFM movie; centralspindlin

As the second example here, we applied the Kalman smoother method to a HS-AFM movie of centralspindlin (29). The cell division contains many phases, one of which divides the cytoplasm into two daughter cells, which is called the cytokinesis. The centralspindlin plays the central roles in this cytokinesis bundling the microtubules and anchoring the plasma membrane within the cytoplasm (Figure 11A top). The centralspindlin consists of two copies of MKLP1 (called ZEN4 for *C. elegans*), which is a motor protein kinesin-6, and two copies of non-motor protein CYK4 (Figure 11A bottom. Note that mAG is not a natural component. See below.).

**Figure 11.**
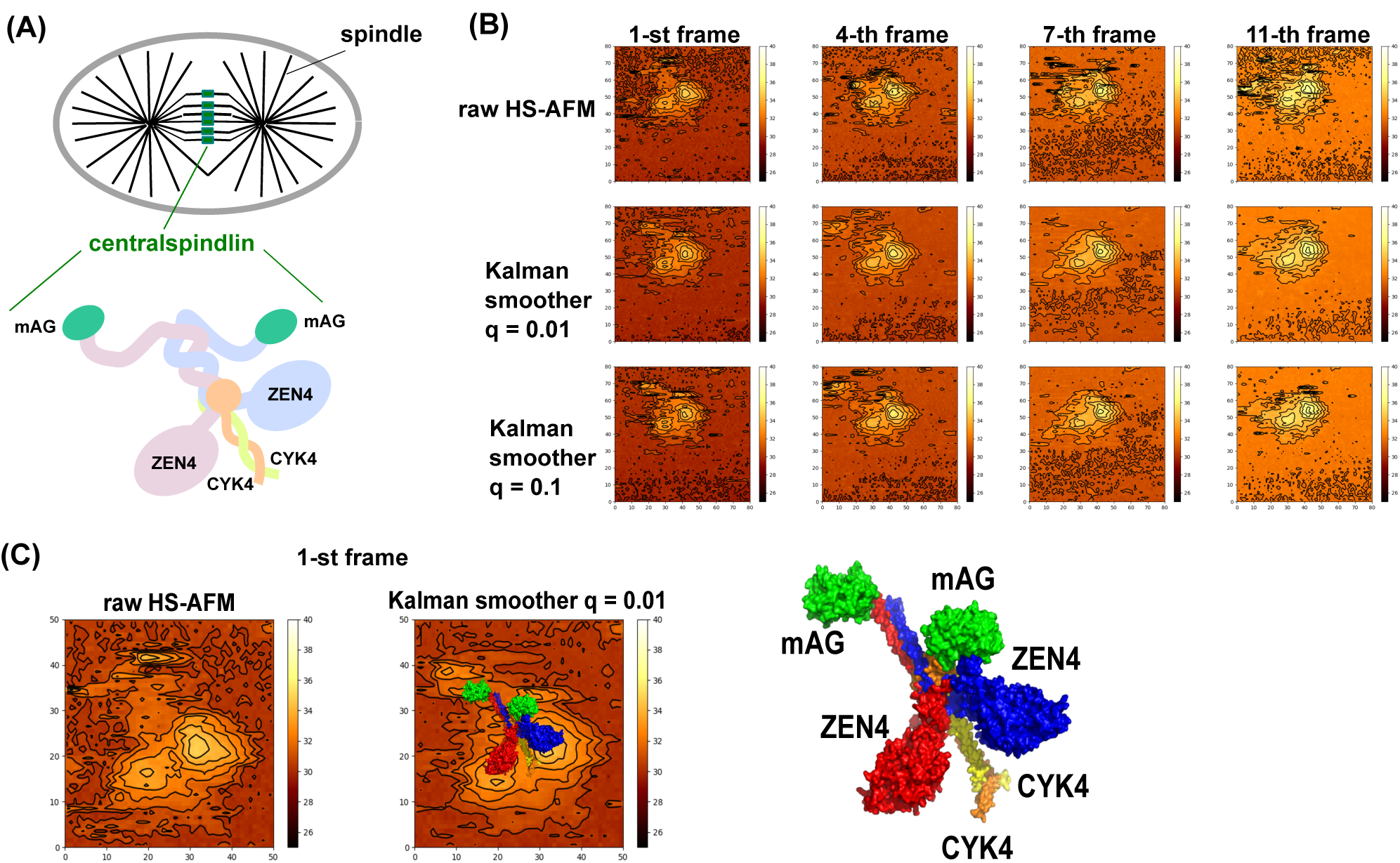
The Kalman smoother for a real HS-AFM movie of centralspindlin. (A) Cytokinesis, a part of cell division, divides the cytoplasm into two daughters’ ones. Centralspindlin, a central molecule in cytokinesis, bundles microtubule and anchors membrane at the emerging boundaries between two daughter cells. Centralspindlin is a tetramer containing two copies of kinesin-family motors MKLP1 (ZEN4) and two copies of RhoGAP factor CYK4. (B) Some frames (1-st, 4-th, 7-th, and 11-th) of raw HS-AFM movie (top), movies obtained by the Kalman smoother with q = 0.01 (middle), and q = 0.1 (bottom) are shown. (C) For the 1-st frame, a modeled conformation of centralspindlin is superimposed on the image by the Kalman smoother (middle). Labels are given in the right picture. mAG is an artificially inserted domain for ease of identification of disordered tails.

Recently, the HS-AFM measurement provided structural dynamics of many different constructs of centralspindlin from *C. elegans* (29), to which we apply the Kalman smoother here. ZEN4 (a kinesin-6 molecule) consists of the kinesin motor domain responsible for ATP-driven motor activity and the coiled-coil regions that are responsible for homo-dimerization and for binding to CYK4, in addition to C-terminal disordered region. To visualize the C-terminal disordered region, a GFP-like domain, called mAG, is artificially added to the C-terminus end that serves as a marker in HS-AFM (Figure 11A bottom).

The raw HS-AFM movie shows, albeit noisy, two globular shapes together with some extrusion visible in some, but not all, frames (Figure 11B, the top row, Movie S10). We applied the Kalman smoother method with two parameter values, q = 0.01 (the middle row, Movie S11) and q = 0.1 (the bottom row) to a short part of a movie containing 13 frames (denoted as 0-th to 12-th frames) of 80×80 pixels. As in the case of FlhAc, the results in Figure 11 show that the Kalman smoother clearly reduced the noise and revealed clearer two globular shapes at the center and some extrusion in the middle of the two globules. Surprisingly, at the initial and the 4-th frames in Figure 11B, we recognize two small spots at the top-left of the image. Looking back the raw HS-AFM images (the top row) of the same frames, we do not find the two small spots, but occasionally see one spot. Possibly, due to the fast diffusion of that part, the data asynchronicity may cause such distortions in the raw HS-AFM images.

Using a relatively well characterized 1-st frame of the Kalman smoother image, we try to map an atomic-resolution structural model. We first performed a standard loop structure modeling for ZEN4, CYK4 and mAG, using the modeling server Phyre2 (30). We excluded a loop region near the neck linker of ZEN4, due to its apparently poor model. We could assign the two large globular shapes of the Kalman smoother image as the two kinesin motor domains, the extrusions as the CYK4, and the two small spots as the engineered mAG domains. Then, with the mapping of 0.7375 nm per pixel, we aligned each molecular structure model on the Kalman smoother image, comparing the Kalman smoother image with the AFM-like image of the model structure created by afmize (the probe tip radius 1nm and the apex angle 10º)(Figure 11C). The AFM-like image from the molecular model captures major features in the Kalman smoother image. Probably, the mAG domain attached at the end of disordered region moves relatively rapidly and the identification of the two small spots (assigned as mAG domains) shows a good performance of the Kalman smoother to resolve the data asynchronicity.

Comparing the two parameter values, the q = 0.01 gives better denoising effect, while q = 0.1 may capture the instantaneous structure change more rapidly.

## Discussion & Conclusion & Future Work

We examined the Kalman filter and smoother methods to resolve the data asynchronicity in HS-AFM data. Twin-experiments for both the diffusing cone model and the conformational change in a motor protein dynein showed that the Kalman smoother method can reduce the data asynchronicity in HS-AFM movie. The Kalman smoother method also serves to reduce the noise. Applications of the Kalman smoother method to two real HS-AFM movies, FlhAc and centralspindlin, produced markedly clearer images than the raw HS-AFM movies, which enables us to construct structural models of the target molecules. The method is general and can be applicable to any HS-AFM movies.

This study is the first trial to apply the data assimilation approaches to the HS-AFM data, which thus has much room of improvement. Especially, we used a very simple system model *F*_*t*_ in this study; the identify matrix for most of the results and a simple diffusing propagator in one case. In general, we can optimize the system matrix by maximizing the likelihood, or the posterior probability. This optimization, however, is computationally not straightforward. As another direction, we detect structural features in the HS-AFM images, from which we can suggest more mechanistic system model for each target molecules.

## Supplemental Movies

**Movie S1**. A ground-truth AFM-like movie of a diffusing cone.

**Movie S2**. A raw AFM-like movie of a diffusing cone.

**Movie S3**. A movie obtained by the Kalman filter method (q=0.1) for a diffusing cone.

**Movie S4**. A movie obtained by the Kalman smoother method (q=0.1) for a diffusing cone.

**Movie S5**. A ground-truth AFM-like movie of a dynein conformational change dynamics

**Movie S6**. A raw AFM-like movie of a dynein conformational change dynamics

**Movie S7**. A movie obtained by the Kalman smoother method (q=0.01) for a dynein conformational change dynamics

**Movie S8**. A HS-AFM of FlhAc

**Movie S9**. A movie obtained by the Kalman smoother method (q=0.01) for FlhAc

**Movie S10**. A HS-AFM of centralspindlin

**Movie S9**. A movie obtained by the Kalman smoother method (q=0.01) for centralspindlin

